# Condensins-mediated mitotic folding persists as a structural echo in interphase chromatin

**DOI:** 10.1101/2025.08.08.669285

**Authors:** Anastasia Yunusova, Egor Tsoi, Alexander Smirnov, Tatiana Shnaider, Inna Pristyazhnuk, Maria Gridina, Evgeny Kazakov, Igor Kireev, Myakinkov Ivan, Miroslav Nuriddinov, Veniamin Fishman, Nariman Battulin

## Abstract

The Structural Maintenance of Chromosomes (SMC) protein complexes, cohesin and condensins, orchestrate cell cycle-dependent transitions in chromosome folding: from the relatively decondensed interphase chromatin, organized into dynamic DNA loops by cohesin, to the highly condensed arrays of DNA loops in mitotic chromosomes formed by condensins. Here, using Hi-C data from mouse embryonic stem cell lines with auxin-inducible degron-tagged subunits of SMC complexes, we clarify the role of condensins in shaping chromatin contacts within the interphase nucleus. We found that depletion of condensin I leads to weakened segregation of chromatin into A and B compartments. We also show that the contact patterns established by condensins during mitosis are preserved in interphase, suggesting a structural memory of mitotic chromosome folding. Condensins influence chromocenter organization: in the absence of condensin II during mitosis, centromeres tend to cluster, forming hyperclusters of pericentric heterochromatin. In contrast, depletion of condensin I reduces the number of heterochromatin clusters, consistent with enhanced chromosome territoriality. By combining a cohesin-degron system with Mcph1 knockout, we established an artificial model in which condensin II becomes the major driver of chromatin folding in interphase nuclei following cohesin depletion. Unlike extrusive cohesin, condensin II does not stall at sequence-specific sites and forms large, randomly positioned loops on interphase chromatin. Moreover, condensin II activity during interphase does not interfere with cohesin-dependent structures such as TADs and chromatin loops.

**Graphical abstract:** 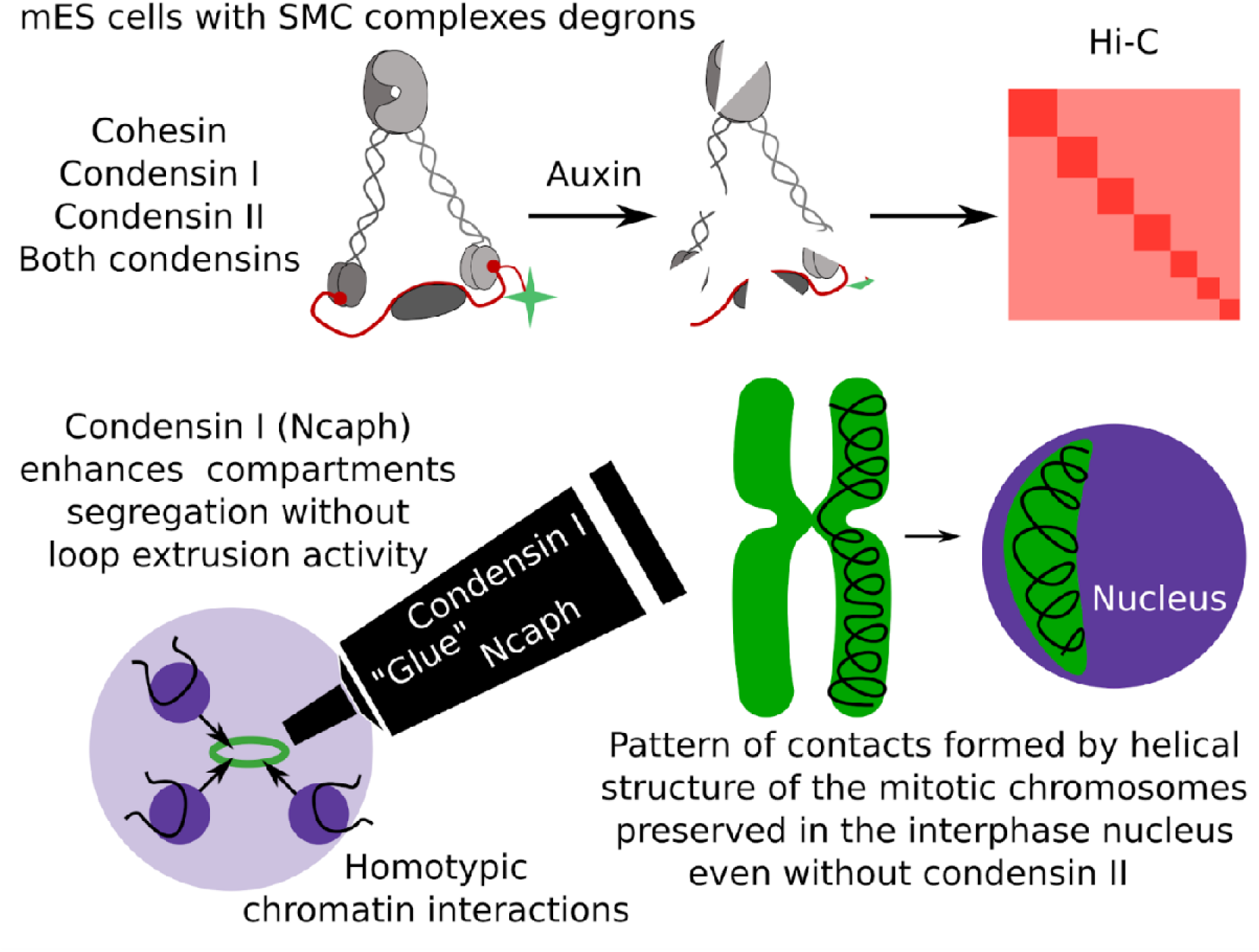

## INTRODUCTION

Chromatin architecture dynamically transforms through the cell cycle, from compact metaphase chromosomes to more relaxed interphase chromatin, where active and repressed sequences co-segregate into distinct spatial compartments (Yatskevich et al. 2019). Cohesin and condensins (I and II), members of the SMC protein complex family, are key mediators of chromatin organization, playing essential roles in mitotic chromatin compaction, transcriptional regulation, and DNA repair (Hoencamp and Rowland 2023).

The cohesin core complex is composed of SMC1, SMC3, and non-SMC subunits: the kleisin family protein RAD21 and the chromatin-associated proteins SA1 or SA2 (Larionov et al. 1985; Michaelis et al. 1997). Condensins I and II share the same pair of SMC subunits (SMC2 and SMC4), but differ in their sets of non-SMC components: NCAPH, NCAPG, and NCAPD2 for condensin I, and NCAPH2, NCAPG2, and NCAPD3 for condensin II (Figure 1A) (Neuwald and Hirano 2000). Both cohesin and condensins organize large-scale chromosome structures through ATP-dependent loop formation, a process known as chromatin loop extrusion (Fudenberg et al. 2016; Ryzhkova et al. 2024).

**Figure 1.**
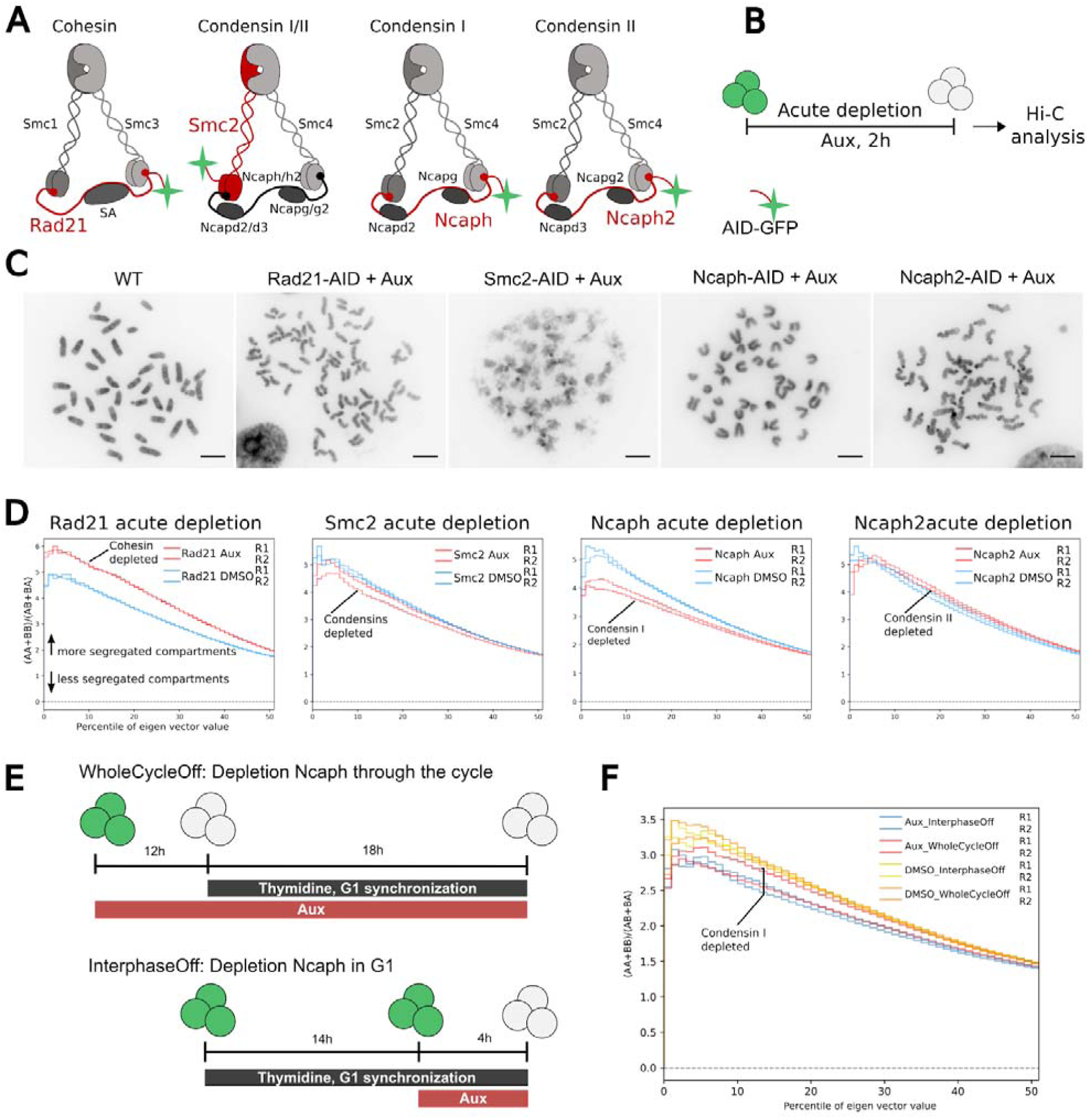
Ncaph depletion during interphase decreases chromatin compartmentalization. **A** - Schematic illustration showing the degron-tagged subunits of SMC protein complexes. The red color represents the cohesin/condensins subunits fused to the AID-GFP tag (marked by green color) and targeted by proteasome degradation after auxin exposure. **B** - Scheme of the experiment. **C** - Representative microscopy images of mitotic chromosome spreads with condensation defects after cohesin or condensins depletion. Scale bar 5 μm. **D** - Saddle strength plots for the whole genome following cohesin (Rad21-AID), both condensins (Smc2-AID), condensin I (Ncaph-AID), and condensin II (Ncaph2-AID) depletion in two biological replicates. **E** - Scheme of the experiment with cell arrest at G1 by thymidine treatment. **F** - Saddle strength plots for the whole genome following Ncaph depletion through the cycle and during G1 only.

Throughout interphase, cohesin extrudes genomic DNA into loops until it encounters barrier proteins or is removed from chromatin by WAPL (Rao et al. 2014). Thus, cohesin mediates chromatin contacts between distant genomic sites, promoting gene regulation, DNA repair, and recombination events. In population-averaged Hi-C maps, positions where cohesin is stalled appear as discrete dots and triangle-shaped structures along the diagonal, known as topologically associating domains (TADs) (Dixon et al. 2012; Nora et al. 2012). TADs contribute to proper gene expression by promoting or restricting interactions between cis-regulatory elements and their targets (Kabirova et al. 2024). In addition to its role in TAD organization, cohesin also modulates chromatin compartmentalization, which appears on Hi-C maps as a “checkerboard” pattern. The prevailing view is that compartmentalization arises via phase separation driven by homotypic interactions between A (transcriptionally active) and B (transcriptionally repressed) genomic regions (Lieberman-Aiden et al. 2009). It is believed that cohesin-mediated loop extrusion interferes with compartmental interactions, suggesting that TADs and compartments are organized by fundamentally distinct and opposing forces (Mirny et al. 2019).

Condensins play an important role in structuring chromatin within the mitotic chromosome (Hirano and Mitchison 1994; Hirano 2025). Condensin II organizes chromatin into large outer loops, which are further subdivided into smaller nested loops by condensin I. By accumulating along the length of metaphase chromosomes, both complexes ensure proper chromosome condensation and segregation during mitosis and meiosis. The resulting densely packed mitotic chromosomes are folded around a centrally located helical scaffold, resembling a bottlebrush or spiral staircase (Gibcus et al. 2018).

The binding of condensins and cohesin to chromatin is tightly regulated throughout the cell cycle (Hibino et al. 2024; Hildebrand et al. 2024; Samejima et al. 2025). Cohesin remains localized in the nucleus throughout interphase and is released from chromatin at prophase, while a residual pool is retained at the centromeres of sister chromatids until anaphase onset. Condensin II also has nuclear localization but becomes activated and stably associates with chromatin only in early prophase. One of the key factors that prevents unscheduled condensin II activity during interphase is MCPH1. MCPH1 inhibits the binding of condensin II subunits, thereby blocking condensin II loading onto chromatin. Loss of MCPH1 leads to premature chromatin condensation during G1 and G2, resulting in an increased fraction of cells displaying prophase-like chromosome organization (Houlard et al. 2021). Condensin I is loaded onto chromosomes immediately after nuclear envelope breakdown in prometaphase and dissociates near the time of a new nucleus formation. During interphase, condensin I is generally thought to be localized in the cytoplasm. However, there is evidence that a subset of condensin I complexes may remain in the nucleus throughout interphase (Brunner et al. 2025).

Whether condensins perform any function related to the formation or maintenance of chromatin loops in the interphase nucleus remains unclear. Since condensin II is localized in the nucleus, it has been debated whether it contributes to interphase chromatin organization (Wallace et al. 2015; Hoencamp et al. 2021). Some studies have reported that condensin II colocalizes with architectural and insulator proteins at domain boundaries and is also recruited to super-enhancers, suggesting a possible role in gene regulation (Dowen et al. 2013; Li et al. 2015; Hassan et al. 2020; Marshall et al. 2020). Other studies, however, have proposed that changes in gene expression observed after condensin II depletion are indirect consequences of mitotic defects (Hocquet et al. 2018). Furthermore, several reports indicate that a substantial fraction of condensin I remains in the nucleus after cell division (Brunner et al. 2025). How this nuclear pool of condensin I may affect chromatin architecture during interphase is still unknown.

Here, we used an auxin-inducible protein degradation system to determine the contribution of cohesin and condensin subunits to interphase chromatin organization in mouse embryonic stem cell lines. We showed that depletion of condensin I results in weakened segregation of A and B chromatin compartments, an effect that appears to be independent of its loop-extruding activity. Depletion of either condensin I or II alters the mitotic chromosome loop pattern, which in turn influences interphase chromatin contacts—a phenomenon we refer to as the structural echo of the mitotic chromosome. Depletion of condensin II increases chromocenter clustering during interphase, whereas depletion of condensin I has the opposite effect, leading to reduced chromocenter clustering. Overloading of condensin II in interphase, as observed in Mcph1 knockout cells, does not result in the formation of distinct structures similar to those generated by cohesin through loop extrusion. Thus, condensins influence contact formation in the interphase nucleus primarily through their effects on mitotic chromosome architecture. In addition, condensin I plays a significant role in maintaining the segregation of chromatin compartments in the nucleus.

## RESULTS

### Mouse ES cells with inducible depletion of cohesin and condensins

To investigate chromatin interactions mediated by architectural SMC proteins, we selectively inactivated cohesin and condensin subunits using auxin-inducible degron technology. Using CRISPR/Cas9 genome editing, we previously established mouse embryonic stem (ES) cell lines in which the endogenous loci of Rad21 (cohesin), Ncaph (condensin I), Ncaph2 (condensin II), and Smc2 (both condensins) were homozygously tagged with the miniIAA7-mEGFP degron (Figure 1A, Figure S1A) (Yunusova et al. 2021). These engineered cell lines maintained stable phenotypes and exhibited rapid and efficient degradation of degron-tagged proteins upon auxin treatment (Figure S1B, S1C).

Prolonged depletion of Rad21 or Smc2 led to cell cycle arrest and cell death due to defective chromosome segregation (Figure S1D). In contrast, separate depletion of condensin I or condensin II did not produce such severe effects and caused no detectable changes in cell cycle distribution (Figure S1D). The efficiency of SMC complex depletion in these ES cell lines was most clearly demonstrated by altered mitotic chromosome morphology upon auxin treatment. Rad21-depleted cells displayed severe defects in sister chromatid cohesion. In Smc2-depleted cells, mitotic chromosomes failed to condense and appeared as diffuse, cloud-like chromatin. Condensin II promotes axial shortening of chromosomes; accordingly, Ncaph2-depleted cells exhibited elongated and thinner chromatids with a characteristic curly morphology. Conversely, depletion of condensin I, which facilitates lateral compaction, resulted in defective folding with “swollen” chromatids that appeared shorter and thicker (Figure 1C). Thus, this panel of engineered ES cell lines enables us to study the consequences of acute depletion of individual SMC complexes on chromosome architecture.

### Depletion of condensin I decreases chromatin compartmentalization

We treated all cell lines with auxin for 2 hours and then performed Hi-C experiments (Figure 1B; see also Supplementary Table S1 for sequencing and quality control metrics). Cohesin depletion caused a strong reduction in TADs and chromatin loops in mouse ES cells, consistent with previous reports. In contrast, depletion of condensins—either condensin I or II individually, or both simultaneously—did not result in significant changes in TADs or loops (Figure S2). These structures are known to be formed by cohesin complexes stalled at CTCF binding sites.

We next evaluated the effects of acute depletion of SMC complexes on chromatin compartmentalization. Consistent with previous studies, we observed that cohesin-mediated loop extrusion antagonizes compartmentalization. As expected, cohesin depletion led to an increase in compartmentalization (Figure 1D). Depletion of condensin II, or both condensins simultaneously, did not result in major changes in compartmentalization (Figure 1D). Interestingly, depletion of condensin I resulted in a significant decrease in compartmentalization (Figure 1D). Notably, the magnitude of this effect was comparable to that observed with cohesin depletion—but the direction of change was opposite, leading to a weakening of compartmentalization.

Since we used an asynchronous cell population, we could not determine whether the observed decrease in compartmentalization results from defects in chromatin condensation during mitosis or occurs directly during interphase. To resolve this, we arrest cells at the G1/S transition using thymidine (Figure S3). We designed two experiments: (1) auxin was added 12 hours before thymidine treatment, resulting in condensin I depletion throughout the entire cell cycle (WholeCycleOff) (Figure 1E); (2) auxin was added 14 hours after thymidine treatment, when cells were already arrested at the G1/S boundary, so that condensin I was present during mitosis but depleted during interphase (InterphaseOff). In both conditions, we observed a significant decrease in compartmentalization (Figure 1F). Thus, even during interphase, Ncaph depletion leads to a notable reduction in A/B compartmentalization.

### The shape of the mitotic chromosome sculpted by condensin II is preserved throughout interphase

To further analyze Hi-C contact maps after cohesin and condensin depletion, we computed contact probability curves, P(s), and their slopes. P(s) reflects the probability of contact between loci separated by a given genomic distance. The proximal region of the curve, spanning approximately 1 Mb (referred to as the “shoulder”), captures the local enrichment of contacts driven by cohesin-mediated loop extrusion—one of the key features of interphase chromatin organization. The distal region of the P(s) curve reveals chromosome architecture at the megabase scale (5–10 Mb), with its most prominent feature being the “second diagonal,” which represents the helical scaffold that forms during mitotic chromosome condensation (Figure 2A-F). The slope of P(s) is particularly informative for detecting subtle changes in contact probabilities. It is important to note that P(s) is derived from a whole-genome Hi-C dataset, so even small shifts in the curve’s shape can reflect changes in the underlying principles of chromatin contact organization.

**Figure 2.**
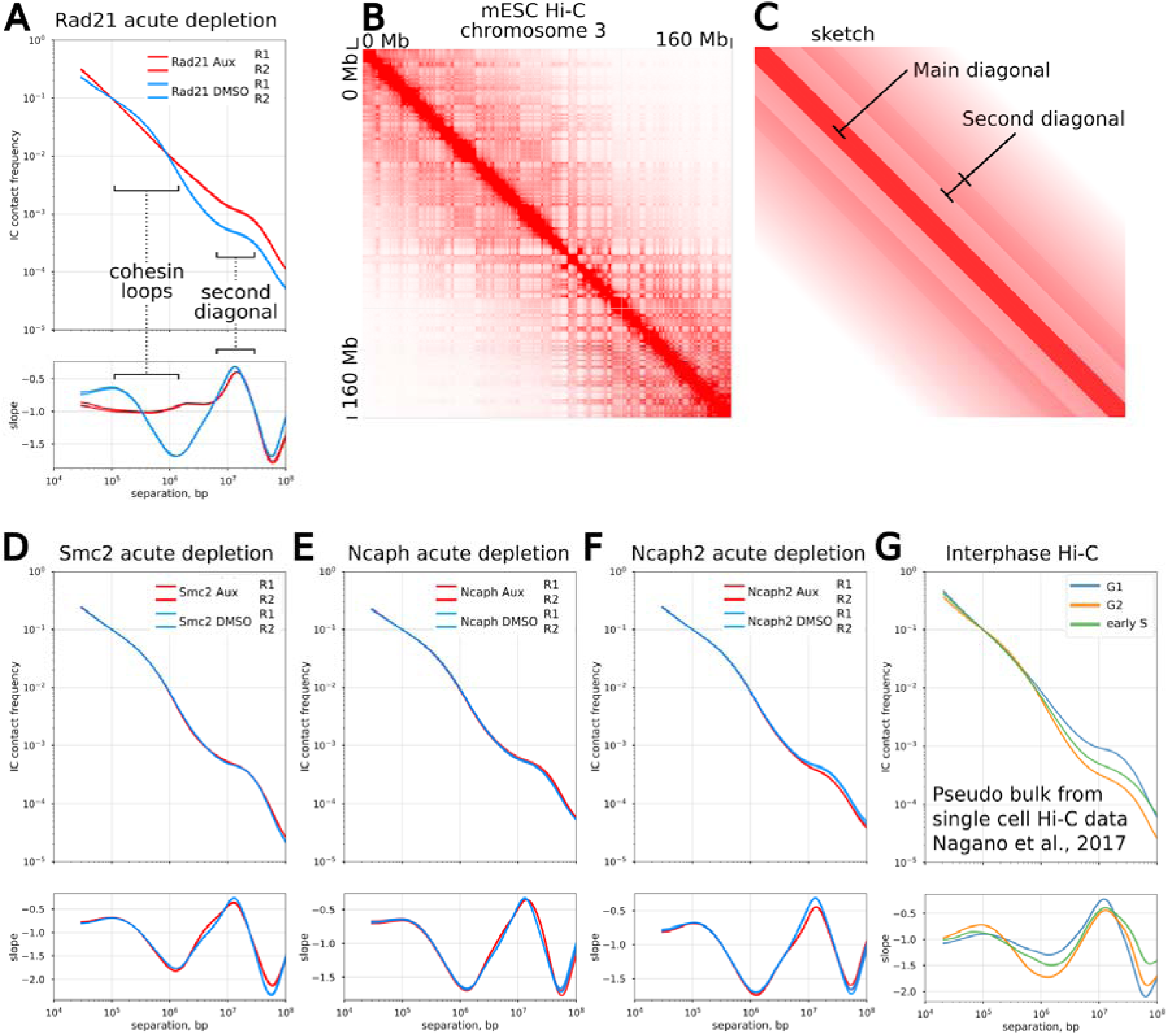
The second diagonal band reflecting the mitotic chromatin folding persists during interphase/ Hi-C contact maps upon acute cohesin and condensins loss. **A** - Chromosome-averaged distance-dependent contact probability P(s) and slopes in control and after cohesin (Rad21-AID) depletion in two biological replicates. **B** - Hi-C contact maps and its sketch (C) of mouse chromosome 3 at 10-Kb resolution illustrating the second diagonal band. (**D-F**) - Chromosome-averaged distance-dependent contact probability P(s) and slopes in control and both condensins (D), condensin I (E), and condensin II (F) depletion in two biological replicates. **G** - P(s) and slopes from single-cell Hi-C data distributed by cell cycle stages.

Since cohesin depletion disrupts TADs, the shoulder of the P(s) curve in Rad21-depleted cells displayed a significantly shallower slope (Figure 2A). In contrast, the shoulders of P(s) in Ncaph-, Ncaph2-, and Smc2-depleted cells were indistinguishable from those of untreated controls (Figure 2D–F). Thus, at the level of TADs, depletion of condensins does not lead to detectable changes in chromatin contacts.

The most prominent changes in contact patterns were observed at ∼15 Mb genomic separations, corresponding to the second diagonal. Following depletion of condensin II or both condensins, only a slight reduction in the 15 Mb second diagonal peak was detected (Figure 2D, F). It was previously shown that the second diagonal band arises during prometaphase and is mediated entirely by condensin II (Gibcus et al. 2018). Based on the assumption that the 15 Mb peak reflects a mitosis-specific feature, we expected this signal to be absent in condensin II-depleted cells compared to untreated controls (Figure 2F).

However, the persistence of this signal prompted us to consider whether the second diagonal—a hallmark of mitotic chromatin folding—might also be retained in interphase cells as a “memory of mitotic folding.” Since our experiments used asynchronous cell populations, we could not rule out the contribution of mitotic cells to the observed P(s) profiles. To address this, we turned to single-cell Hi-C data from mouse ES cells sorted by cell cycle stage (Nagano et al. 2017). We generated pseudo-bulk Hi-C maps for G1, early S, and G2 phases, and computed P(s) curves. Strikingly, we observed a prominent second diagonal band in all interphase stages, indicating that this periodicity of interactions—originally established during mitosis—persists throughout interphase (Figure 2G). Furthermore, the continued presence of the second diagonal after acute Ncaph2 depletion suggests that condensin II is not required to maintain this mitotic helical architecture in interphase.

### Depletion of condensin II throughout mitosis leads to altered chromatin structure in the subsequent interphase

To determine whether prolonged loss of condensins results in additional changes in chromatin architecture, we repeated the same set of analyses using Ncaph- and Ncaph2-degron mouse ES cell lines, treating them with auxin for 24 hours (Figure 3A). Since ES cells have a short cell cycle duration (∼12 hours), every cell in the population is expected to progress through mitosis at least once during this time frame in the absence of condensins.

**Figure 3.**
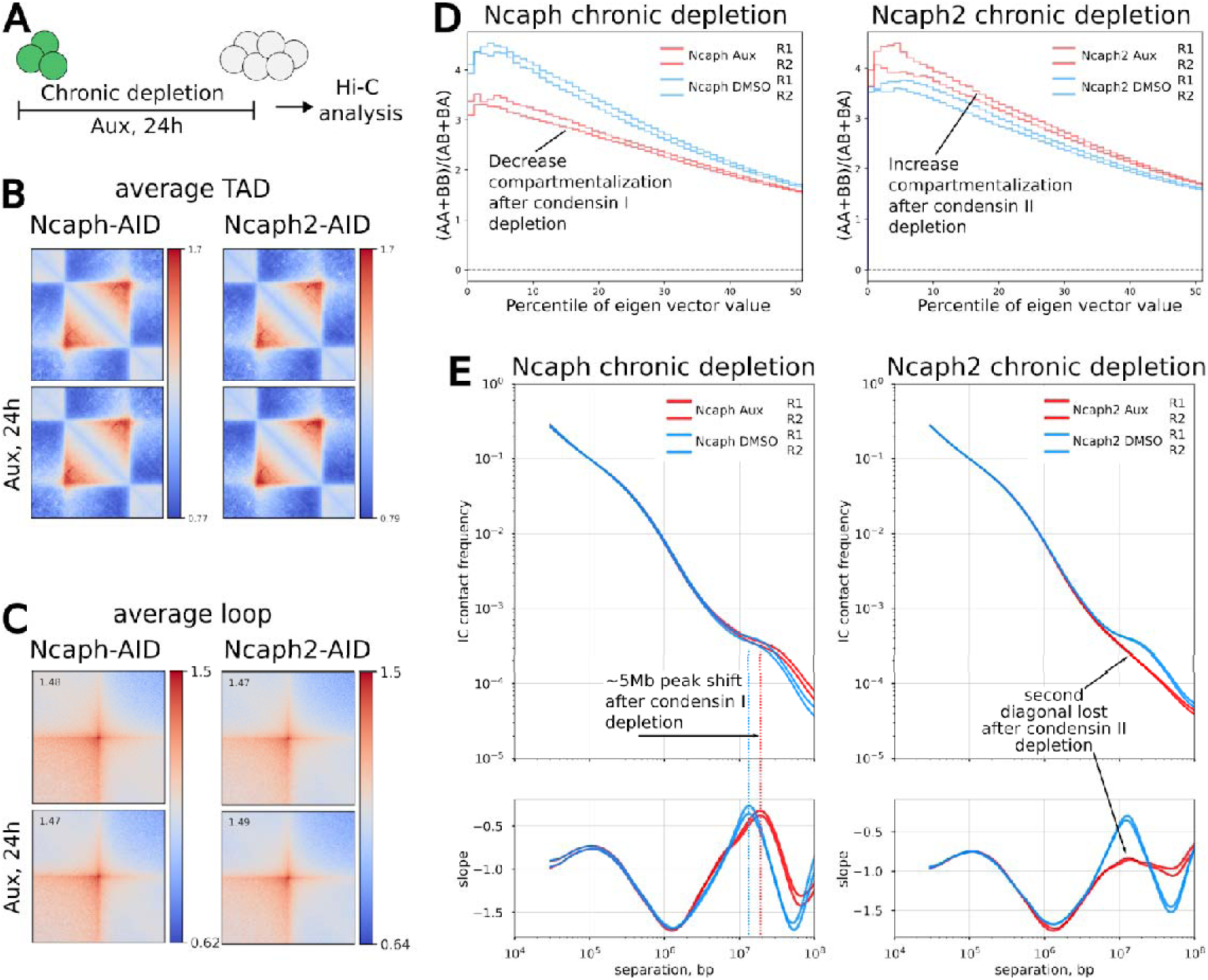
Genome organization changes upon the condensins loss through the cell cycle. **A** - Scheme of the experiment. (**B, C**) - Aggregate TAD (B) and aggregate loop (C) analyses of Hi-C maps in cells depleted condensin I (Ncaph-AID) and condensin II (Ncaph2-AID) for 24h. **D** - Saddle strength plots for the whole genome following condensin I (Ncaph-AID) and condensin II (Ncaph2-AID) depletion in two biological replicates. **E** - Chromosome-averaged distance-dependent contact probability P(s) and slopes in control and after condensin I (Ncaph-AID) and condensin II (Ncaph2-AID) depletion in two biological replicates. A red dotted line indicates the highest frequency of contacts in the second diagonal.

Aggregate TAD and aggregate peak analyses revealed no significant differences in the average Hi-C signal between treated and untreated degron cell lines (Figure 3B,C). We then proceeded to analyze A/B compartmentalization in cells after chronic depletion of condensin I and II. The compartment profiles showed alterations similar to those observed after acute depletion. Specifically, there was a slight increase in compartment strength in the absence of condensin II, and a much more pronounced decrease in the absence of condensin I (Figure 3D).

The most striking observation comes from comparing the contact frequency P(s) curves. In chronically condensin I-depleted cells, the second diagonal appears at a greater genomic distance compared to control cells. We interpret this shift as an extension of outer chromatin loops mediated by the remaining condensin II, along with the persistence of the spiral scaffold characteristic of mitotic chromosomes (Figure 3E). In contrast, in the P(s) curves for cells lacking NcapH2, we observed a complete absence of the second diagonal band compared to untreated cells (Figure 3E). This is consistent with previous findings showing that NcapH2 depletion leads to the disappearance of the second diagonal band in P(s) curves during mitosis (Gibcus et al., 2018). Given our results - specifically, that the second diagonal band is present in interphase P(s) curves and remains unchanged after acute condensin II depletion - this supports the idea that the contact pattern formed by the helical twisting of loops in mitotic chromosomes is retained into the subsequent interphase.

### Condensin II does not stall at sequence-specific sites and does not form distinctive chromatin structures in interphase after cohesin depletion

The small increase in compartmentalization observed upon chronic condensin II depletion may suggest that a subset of condensin II complexes remains associated with chromatin during interphase. To investigate the role of condensin II in interphase more directly, we analyzed cells with condensin II overloading on chromatin. For this purpose, we performed Hi-C analysis on Rad21 auxin-inducible degron with Mcph1-knockout mouse ES cells, as previously described (Yunusova et al. 2024b) (Figure 4A). It is known that mutations in MCPH1 lead to premature condensin II binding to interphase chromatin, resulting in increased chromatin compaction and the acquisition of prophase-like, rod-shaped chromosomes within the nucleus (Houlard et al. 2021). In this context, where two types of loop-extruding factors—cohesin and condensin II—are present simultaneously, it becomes particularly intriguing to explore how these complexes interact and influence each other during interphase.

**Figure 4.**
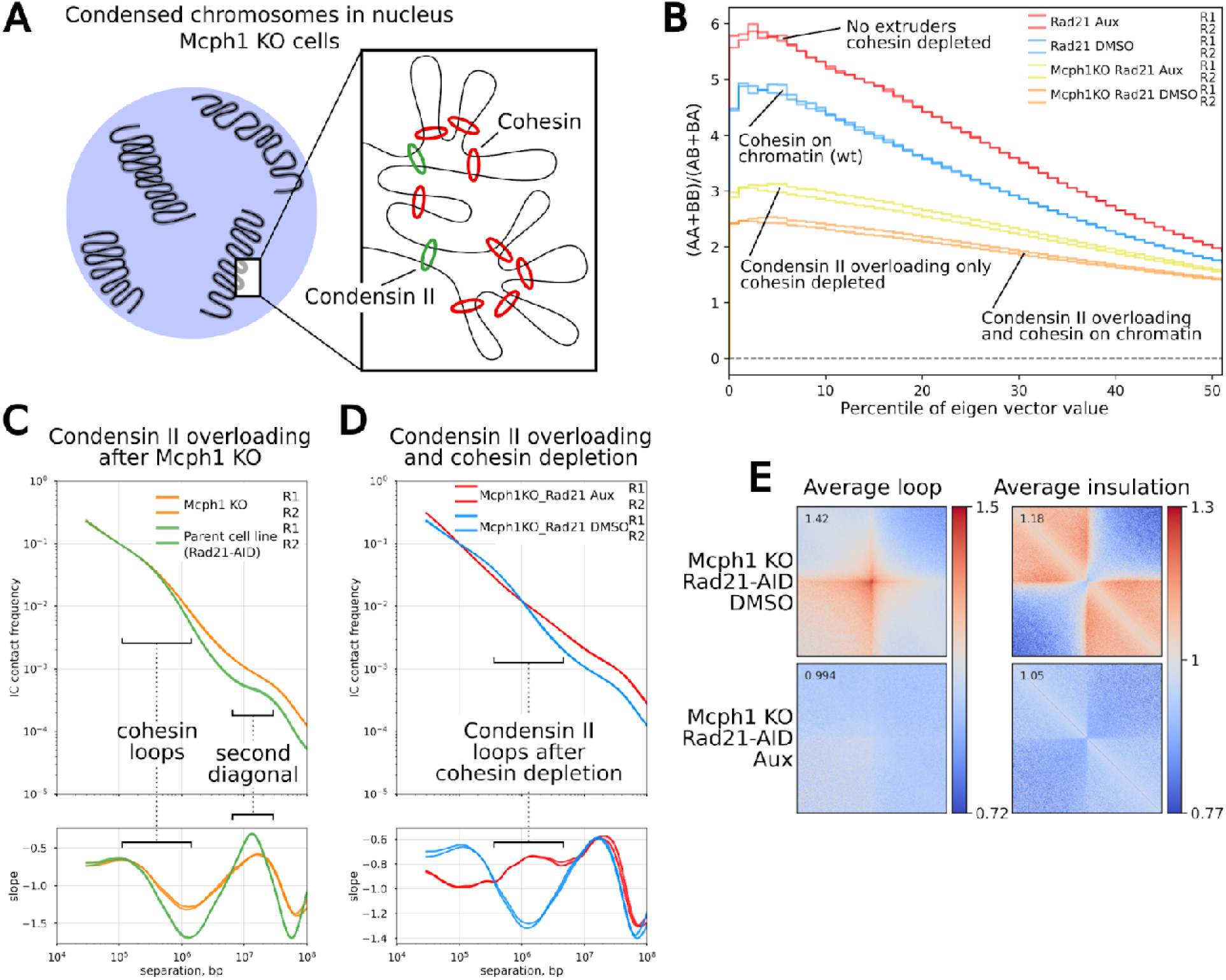
Alterations in chromatin folding caused by condensin II overloading during interphase in Mcph1-KO cells. **A** - Schematic illustration for the formation of condensed prophase-like chromosomes in the nucleus of Mcph1–KO cells. **B** - Saddle strength plots for the whole genome in the presence of different types of extruders on chromatin. (**C,D**) - Chromosome-averaged distance-dependent contact probability P(s) and slopes in Mcph1-KO cells compared to parental line (C) and in Mcph1-KO cells following cohesin (Rad21-AID) depletion (D). **E** - Aggregate loop analysis and average insulation score of the TAD borders in untreated Mcph1–KO cells and after cohesin (Rad21-AID) depletion by auxin exposure for 2 h.

First, we assessed changes in Hi-C features following Mcph1 knockout. The most striking difference between the Hi-C maps of Mcph1-KO cells and the parental line was a dramatic decrease in compartmentalization (Figure 4B). This observation is consistent with the established notion that the loop extrusion process counteracts compartmentalization: the more extruders are bound to chromatin, the more pronounced the reduction in compartmental signal.

It was previously shown that condensins do not accumulate at specific genomic sites; instead, condensin-organized mitotic loops are randomly positioned along the mitotic chromosome (Gibcus et al. 2018). Consistent with this, direct inspection of Hi-C maps from Mcph1-KO cells revealed no evidence of specific condensin II localization within interphase chromatin. TADs and their associated features, such as loops and insulation at domain boundaries, remained unaltered (Figure S4). To test whether cohesin, which persists in interphase, might mask potential condensin-specific features, we induced Rad21 depletion by auxin treatment for 2 hours and then analyzed chromatin contacts. While this treatment led to a marked disruption of cohesin/CTCF-mediated structures such as TADs and loops, we still did not observe any enrichment of specific chromatin features resembling dots or stripes (Figure 4E). These results suggest that condensin II is not capable of forming sequence-specific chromatin structures in interphase.

Then we investigated global chromatin organization by analyzing the distribution of genomic distance-dependent contact frequency (P(s)). Since Mcph1 knockout leads to the formation of visible chromosomes in the interphase nucleus, it is not surprising that the shape of the P(s) curve in Mcph1-KO cells is significantly different from that of parental ES cell lines, where such structures appear only in prophase. At the level of cohesin loops, the P(s) curves are very similar (Figure 4C). This is consistent with our previous finding that condensin II does not affect cohesin-mediated loops anchored at CTCF sites.

However, at larger genomic distances, Mcph1-KO cells show a slower decay in the P(s) curve, indicating increased contact frequency between loci separated by more than 300 kb (Figure 4C). This is closely related to condensin II loading onto chromatin during interphase. Since condensin II has a longer residence time on chromatin than other SMC complexes, the loops it forms are significantly larger. After cohesin depletion by auxin in Rad21-AID Mcph1-KO cells, only interphase condensin II remains. It is clearly seen that it forms loops of much larger size compared to cohesin (Figure 4C, D). It is important to note that we did not detect any additional structures in the Hi-C maps of auxin-treated Rad21-AID Mcph1-KO cells. This suggests that condensin II does not form sequence-specific structures and is randomly distributed across the chromatin.

Finally, the most prominent alterations caused by forced condensin II loading occurred at the level of the helical chromosome structure. First, in Mcph1-KO cells, the second diagonal shifted to a larger genomic distance compared to parental cells (Figure 4C). This relocation suggests that condensin II–mediated loops become longer, which in turn leads to shortening of the helical axis of chromosomes. This shortening is so pronounced that it can be clearly observed by microscopy: metaphase chromosomes in Mcph1-KO cells are significantly shorter than those in control cells (Yunusova et al. 2024b). Moreover, in Mcph1-KO cells, the second diagonal band appeared significantly more diffuse (Figure 4C). Since the second diagonal is thought to reflect interactions between loci located in adjacent loops within a helical turn, this observation suggests that the regular helical order of mitotic chromosomes is partially disrupted following premature condensin II loading during interphase.

Ultrastructural analysis of chromosomes in Mcph1 knockout cells also reveals dense chromatin bodies resembling prophase chromosomes. Interestingly, these “prophase-like” chromosomes are heterogeneous and contain foci of high density, which are interspersed with looser areas (Figure S5). It can be suggested that the areas with a high density correspond to the accumulation sites of condensin II. While the loose areas probably correspond to chromatin loops. It is also worth noting that morphologically, the fibrils inside such “prophase-like” chromosomes resemble the chromonemas identified in other studies (Kireeva et al. 2004; Deng et al. 2016).

Thus, condensin II, when loaded onto chromatin in interphase, does not generate sequence-specific structures, but it does extrude large loops that affect long-range chromatin interactions—extending over distances greater than 1 Mb.

### The increase in compartmentalization is accompanied by pronounced chromosome territories

At the chromosome scale, two types of interphase chromatin organization have been described in eukaryotic cells: (1) each chromosome occupies a distinct region of the nucleus, consistent with the concept of chromosome territories (CTs); and (2) centromeres and telomeres are clustered at opposite sides of the nucleus in the so-called Rabl configuration. The spatial organization of chromosomes in mouse ES cells is consistent with a Rabl-like configuration (Stevens et al. 2017; Houlard et al. 2021). Microscopically, this is characterized by the presence of chromocenters—clusters of centromeric and pericentromeric heterochromatin that appear as Hoechst-dense foci (Figure 5A). Hi-C contact maps along individual chromosomes also demonstrated elevated frequencies of intra-chromosomal interactions between centromeres and telomeres (Figure 5B).

**Figure 5.**
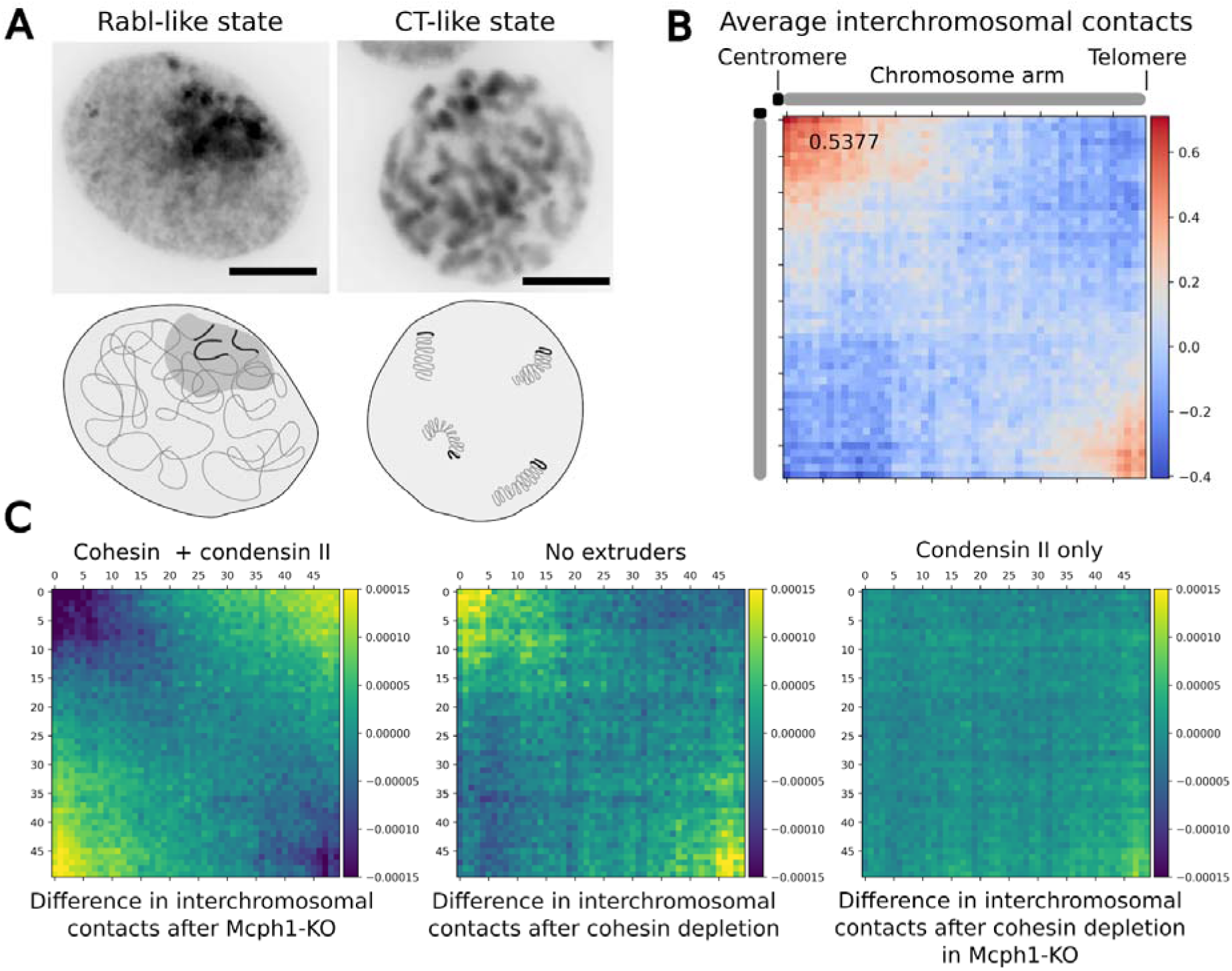
The chromosome configuration of mouse ES cells depends on the amount of extruders in interphase. **A** - Representative images and schematic illustrations showing two types of chromosome configuration: Rabl-like state with centromeres and telomeres clustering together (often observed in mouse ES cells) and CT-like state where chromosomes occupy distinct territories (special case in the interphase nucleus of Mcph1-KO cells). Scale bar: 10 μm. **B** - A typical Hi-C map of average interchromosomal contacts along chromosomes’ length for wild-type mouse ES cells (Rabl heatmap). **C** - Difference in Rabl heatmaps between wild-type cells and cells with different types of chromatin extruders in the interphase nucleus.

We were therefore interested in how changing the amount of chromatin-bound condensins, either by forced loading during interphase (condensin II) or by depletion (condensins I and II), affects the global chromosomal state in mouse ES cells. First, we analyzed differences in inter-chromosomal contact frequencies between Mcph1-KO cells and the parental control line. Since Mcph1-KO promotes chromosome individualization and disrupts centromere clustering, the Rabl-like configuration in Mcph1-KO cells was significantly altered. The “Rabl heatmap” showing the contact difference (Mcph1-KO vs. control) appeared inverted, indicating impaired spatial interaction frequencies between centromeres and telomeres. This result is consistent with a previous report (Houlard et al. 2021) (Figure 5C).

Our set of cell lines with degrons targeting SMC complexes allows us to better characterize the relationship between the Rabl-like configuration, cohesin loop extrusion, and compartmentalization in mammals. We used our Rad21-degron cell line to study this and compared Rabl heatmaps between auxin-treated and untreated control samples. Interestingly, we found that the Rabl-like configuration becomes more pronounced after cohesin depletion, along with the resulting increase in compartmentalization (Figure 5C). This unidirectional change in both compartmentalization and Rabl-like organization supports the idea that the forces driving A/B compartment formation may also contribute to the clustering of centromeres and telomeres that leads to the Rabl configuration.

In contrast, cohesin depletion in Mcph1-KO cells did not lead to any noticeable changes in chromosome organization (Figure 5C). Presumably, in this case, the forced loading of condensin II onto chromatin interferes with centromere and telomere clustering after cohesin depletion.

### Condensin depletion shifts chromosome configurations due to aberrant mitosis-to-interphase transition

Condensin II is thought to determine the global chromosome configuration: chromosome territories or a Rabl-like configuration depending on the presence or absence of condensin II, respectively (Hoencamp et al. 2021). We therefore analyzed chromosome organization following condensin depletion in our Ncaph and Ncaph2 degron cell lines. We did not observe significant changes in Rabl heatmaps after acute condensin depletion (Figure 6A). This fits with the idea that the Rabl configuration is inherited from the anaphase chromosome structure. Simply put, most cells likely did not pass through mitosis and enter the next interphase during the short auxin treatment (2 hours).

**Figure 6.**
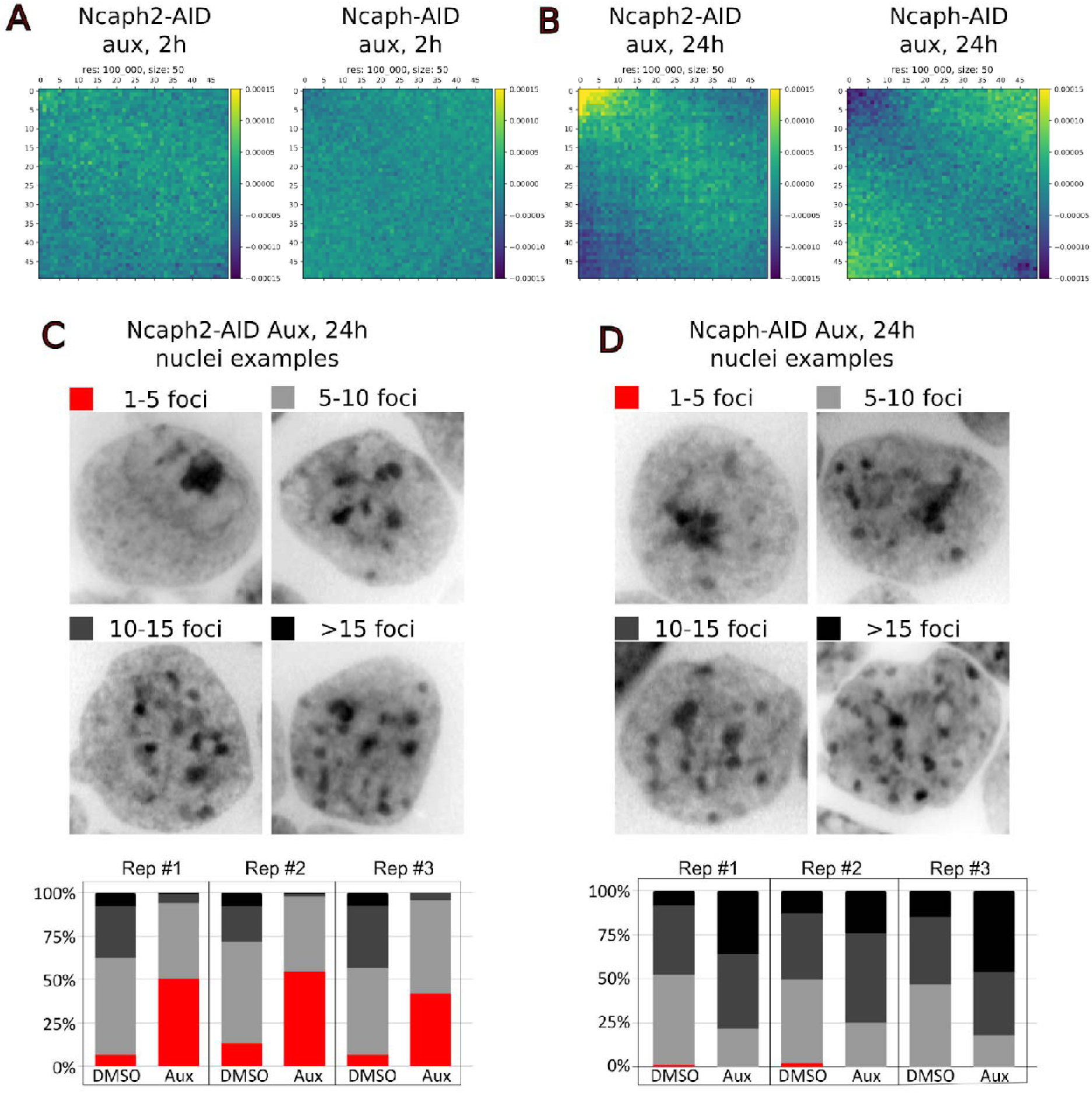
Chromosome configuration changes upon chronic condensins loss. (**A, B**) - Difference in Rabl heatmaps between control cells and cells after acute condensins depletion (A) and chronic condensins depletion (24h). (**C, D**) - Example images of nuclei with different numbers of Hoechst-dense foci in untreated cells and cells depleted of condensin II (Ncaph2-AID) (C) and (Ncaph-AID) (D) for 24h. Quantification of the number of Hoechst-dense foci per nucleus in cells with or without auxin exposure. The experiment was performed in three biological replicates (three independent degron lines for each condensin), with total cell counts of n = 513 (Ncaph2-DMSO), 601 (Ncaph2-Aux), 429 (Ncaph-DMSO), and 384 (Ncaph-Aux). A chi-square test revealed a significant association between au in treatment and the number of Hoechst-dense foci per nucleus across all replicates (p < 0.001).

In the absence of condensin II throughout all phases of the cell cycle, contact frequencies between centromeres were significantly increased (Figure 6B). Consistent with this, we observed a higher proportion of cells displaying hyperclustering of Hoechst-dense foci (Figure 6C). Surprisingly, chronic depletion of condensin I also altered global chromatin architecture, but differently from condensin II depletion. In Ncaph-depleted cells, the Rabl configuration became less pronounced. However, this change was not as substantial as the reduction in centromere and telomere clustering seen in Mcph1-KO cells (Figure 5C). This suggests that chromosome territories are reinforced when cells pass through mitosis without condensin I and enter the next interphase. We hypothesized that these changes would be reflected in heterochromatin organization. Indeed, we observed a decrease in the number of Hoechst-dense foci and an almost complete absence of nuclei with heterochromatin hyperclustering in chronically Ncaph-depleted cells (Figure 6D).

Then we assessed how chronic depletion of Ncaph and Ncaph2 affects the cellular transcriptome. We performed RNA-seq and analyzed gene expression changes after 24 hours of auxin treatment. Although a small number of genes were differentially expressed in each condition (164 genes in Ncaph-depleted cells and 127 genes in Ncaph2-depleted cells), the gene lists overlapped significantly and showed similar expression changes (Pearson correlation coefficient r = 0.95) (Figure S6). We believe that most of these changes are due to activation of the AhR pathway, as previously reported by us and others (Sathyan et al. 2019; Yunusova et al. 2024a). Thus, depletion of Ncaph and Ncaph2 for 24 hours does not lead to significant changes in gene expression.

## DISCUSSION

### Depletion of condensin I results in decreased compartmentalization

The separation of active and inactive chromatin in the nucleus is one of the defining features of interphase chromatin organization. This separation occurs due to the tendency of chromatin regions in a similar epigenetic state to interact preferentially. Although the exact mechanisms behind this homotypic adhesion are not fully understood, microphase separation is believed to play a major role (Belaghzal et al. 2021; Fujishiro et al. 2025). This process is distinct from loop extrusion. In fact, active loop extrusion counteracts chromatin compartmentalization. For example, it has been shown that depletion of cohesin leads to increased compartmentalization (Rao et al. 2017). This likely occurs because removing the loop extruder increases local chromatin mobility (Mach et al. 2022). Without cohesin, the “clips” that prevent chromatin from spreading into homotypic domains are lost, allowing stronger segregation. This logic also explains the increase in compartmentalization we observe after cohesin depletion, as well as the decrease in compartment strength in Mcph1-KO cells, where excess condensin II is loaded onto chromatin. However, we observe an unexpected result in cells depleted of condensin I: a decrease in compartmentalization.

Condensin I is the most abundant SMC complex in cells, but current models suggest that it functions mainly during mitosis to shape metaphase chromosome loops. It was long thought that condensin I is excluded from the nucleus during interphase. However, recent studies show that up to half of the condensin I complexes remain in the nucleus after nuclear envelope reformation (Brunner et al. 2025). Still, there is no clear evidence that condensin I performs loop extrusion during interphase. We therefore speculate that the observed decrease in compartmentalization after Ncaph depletion may not be related to the loss of loop extrusion activity. If condensin I acted purely as an extruder, its removal should increase chromatin mobility and thereby enhance compartmentalization. Yet we observe the opposite. This suggests that condensin I might instead play a role in promoting homotypic chromatin adhesion, and its depletion reduces the effectiveness of this process. A similar reduction in compartment A expression was reported after depletion of BRD2 (Xie et al. 2022). In plants, mutations that alter DNA methylation and histone composition also lead to weakened compartmentalization (He et al. 2024). It is possible that Ncaph has a related role, which may not be directly tied to condensin I function. This hypothesis is indirectly supported by the fact that depletion of another condensin I subunit, Smc2, does not lead to reduced compartmentalization. This suggests that the effect may be specific to Ncaph rather than general to condensin I.

Although Ncaph-depleted ES cells represent an artificial model, it is notable that some human cell types naturally exhibit very low NCAPH expression—such as brain neurons. Interestingly, one of the most striking features of the neuronal 3D genome is weak compartmentalization (Pletenev et al. 2024). Thus, the effect of condensin I on compartmentalization that we observed may have broader biological relevance for chromatin organization in differentiated tissues.

### Condensin II in Interphase

Condensin II is localized in the nucleus during interphase. However, FRAP experiments have shown that under normal conditions, it does not stably bind to chromatin. MCPH1 plays a key role in preventing condensin II loading during interphase, and Mcph1 knockout leads to premature chromatin condensation (Houlard et al. 2021). By combining Mcph1 knockout with Rad21 degradation, we created an artificial system in which condensin II replaces cohesin as the dominant chromatin-bound SMC complex during interphase. This model allowed us to characterize the behavior of condensin II as a potential interphase chromatin extruder. Our results suggest that, unlike cohesin, condensin II does not pause at CTCF sites during interphase. More broadly, we did not observe evidence that condensin II forms sequence-specific structures like those generated by cohesin at defined boundary elements. However, we note a key limitation of our study: we did not directly establish whether condensin II, when loaded due to Mcph1 loss, actively extrudes DNA in interphase. This question remains technically challenging. Even for cohesin and condensins under normal conditions, loop extrusion activity in vivo is inferred from indirect evidence, such as the appearance of characteristic Hi-C patterns (e.g., stripes, fountains) (Vian et al. 2018; Galitsyna et al. 2023) and from polymer modeling studies (Corsi et al. 2023).

Moreover, despite compelling in vitro evidence that both cohesin and condensins can perform loop extrusion, non-extrusion mechanisms of SMC-mediated chromatin folding are still actively discussed (Forte et al. 2025). It is also known that condensins undergo post-translational modifications that vary across the cell cycle (reviewed in (Dekker and Dekker 2022)), so it is possible that condensin II loaded during interphase lacks extrusion activity. Alternatively, it may be passively repositioned by other chromatin-associated factors, such as RNA polymerase or cohesin. Such alternative mechanisms could be further explored using polymer modeling.

### Condensins influence interphase chromatin architecture through mitotic chromosome structure

Our data from condensin-depleted cells suggest that condensins do not play a direct role in loop extrusion in interphase chromatin. However, they clearly influence the global architecture of the interphase nucleus by shaping the structure of mitotic chromosomes, which serve as a template for nuclear organization after cell division. This effect may be described as a structural echo—a residual imprint of mitotic chromosome folding that persists into interphase. This echo is likely most evident in rapidly dividing cells such as embryonic stem cells, as well as in erythroblasts, where the final divisions before terminal differentiation occur rapidly. In addition, erythroblasts exhibit very low transcriptional activity, which may help preserve features of mitotic chromosome organization. We previously reported that vertebrate erythroblasts are characterized by a visible second diagonal on Hi-C maps (Fishman et al. 2019; Ryzhkova et al. 2021). Similarly, a pronounced second diagonal is also present in fast-dividing neural progenitor cells collected in the G0–G1 phase. In contrast, this feature is almost absent in differentiated cortical neurons (Bonev et al. 2017).

Depletion of condensins during mitosis led to observable changes in interphase chromatin organization. In the absence of condensin II, centromeres showed increased clustering, consistent with features of a Rabl-like configuration. Conversely, depletion of condensin I promoted the emergence of more individualized chromosomes and enhanced the formation of chromosome territories. Importantly, we did not observe a strong correlation between these large-scale configurational changes and gene expression changes following chronic condensin depletion in mouse ES cells. This suggests that global 3D organization and transcriptional output can be partially uncoupled, at least over short timescales. Or perhaps even suggests that such global changes in 3D chromatin structure are not important in themselves, but are merely a consequence of other physical constraints and mechanical forces (Solovei and Mirny 2024).

We propose that future studies exploring the role of SMC loop extruders—particularly condensins—in shaping chromatin architecture during development and differentiation will be of significant interest. Understanding how mitotic chromosome structure influences interphase organization may help clarify fundamental principles of genome folding in diverse cell types.

## CONCLUSION

The primary forces shaping 3D genome architecture during interphase are cohesin-mediated loop extrusion, counterbalanced by homotypic chromatin interactions that segregate the genome into A and B compartments. In contrast, mitotic chromosome organization is governed by condensin-dependent loop formation. The effects of SMC complex depletion or overloading that we observed are consistent with current models of chromatin folding. Our study introduces a new dimension to the understanding of interphase chromatin organization—namely, the influence of structural configurations inherited from the preceding mitosis. This mitotic “echo” appears to persist into interphase and contributes to shaping nuclear architecture. One observation that remains unexplained by existing 3D genome models is the reduction in compartmentalization upon Ncaph depletion. This finding suggests that, while the primary mechanisms governing interphase chromatin structure are increasingly well understood, important aspects of genome organization remain to be discovered.

## MATERIAL AND METHODS

### Cell lines and culture conditions

Establishment of degron mouse ES lines has been reported earlier (Menzorov et al. 2019; Yunusova et al. 2021, 2024b). Specifically, here we used ES cell lines with degron-tagged the indicated subunits: NcapH for condensin I, NcapH2 for condensin II, Smc2 for both condensins, and Rad21 for cohesin. One of the Rad21-degron cell lines was used as the parental line for generating Mcph1-knockout cells. All cell lines were grown at 37°C with 5% CO2 and passed every 2 days. Mouse ES cells were cultured on a 0.1% gelatin surface under 2i conditions, which ensures the pluripotency by specifically blocking the MAPK–ERK pathway (PD0325901, 1μM) and glycogen synthase kinase 3 (CHIR99021, 3 μM). The culture media were based on DMEM (Thermo Fisher), supplemented with 7.5% ES FBS (Gibco), 7.5% KSR (Gibco), 1 mM L-glutamine (Sigma), 1% NEAA (Capricorn Scientific, Germany), 0.1 mM β[mercaptoethanol, LIF (1000 U/ml, Polygen), and 1% penicillin/streptomycin (Capricorn Scientific, Germany). 3-Indoleacetic acid (IAA, auxin) (Merck, I2886) was dissolved in NaOH (1M) solution to a concentration of 500 mM, aliquoted, stored at -20°C, and used immediately after thawing. For all experiments, the cells were incubated with 500 μM indole-3-acetic acid (IAA) (Sigma) for the indicated times. For the cell cycle arrest at the G1 phase, cells were treated with thymidine (2mM) for the indicated times.

### Cell Cycle Analysis

Cells were collected, washed with PBS, and then fixed in ice-cold 70% ethanol overnight at 4°C. Prior to cell cycle analysis, cells were centrifuged and stained with a propidium iodide staining solution (500 ug/ml, 100 ug/ml RNase A and 0,1% Triton X-100) for 30 min at RT. Fluorescence detection was performed using an analytical flow cytometer BD FACSAria (BD Biosciences).

### Chromosome spreading

Cells were exposed to auxin for 4 hours and then detached with 0.05% Trypsin-EDTA solution (Capricorn Scientific GmbH, Germany). Then a hypotonic solution (0.25% KCl and 0,2% Sodium citrate) was added to the culture dish for 20 min at 37 °С. Then, cells were collected and fixed with Carnoy fixative (3:1 methanol: glacial acetic acid), dropped onto cold wet glass slides, and stained with 1 μg/ml 4′,6-diamidino-2-phenylindole (DAPI) (Sigma-Aldrich). The samples were analyzed using a Carl Zeiss Axioscop 2 fluorescence microscope. Image processing was carried out using ISIS software (MetaSystems GmbH, Germany).

### Quantifications of Hoechst-dense foci per nucleus

Cells with Ncaph- and Ncaph2 degrons were treated with auxin or left untreated (control) for 24 hours and then collected, washed with PBS, and subjected to cytospin slides preparation at 1400 rpm for 7 min. After fixation with 4% paraformaldehyde in PBS for 10 min at room temperature, slides were rinsed three times and stained with Hoechst 33258 (Sigma-Aldrich). Then slides were rinsed with PBS three times, thoroughly dried, and covered with Mounting medium (Servicebio Technology). Images were acquired by using a Carl Zeiss Axioscop 2 fluorescence microscope with a 63× oil immersion objective lens and encoded for blind analysis using ImageJ software.

### Electron microscopy

Mcph1-KO cells were grown on fibronectin-coated cover glasses and fixed with 2.5% glutaraldehyde solution in 100 mM sodium cacodylate for 2h. Then, samples rinsed three times for 5 min in 100 mM sodium cacodylate, and postfixed in 1% osmium tetroxide in 100 mM sodium cacodylate for overnight at +4 °C. Samples were dehydrated through increasing concentrations of ethanol (50%, 70%, 80%, 96%). Subsequently, mixtures of epoxy resin and ethanol were used with an increase in the resin content. After replacing the mixture with pure resin Epon 812 resin was polymerized at +60°C for 48 hours. Sections of 80 - 90 nm thickness were prepared using a Reichert-Jung Ultracut E ultramicrotome equipped with an Ultra 45 diamond knife. Sections were mounted on formvar-coated copper slot grids and post-stained with 5% aqueous uranyl acetate for 30 min and with lead citrate for 45–90 sec. Sections were examined with a JEM 1400 transmission electron microscope (JEOL, Japan) equipped with a QUEMESA bottom-mounted CCD-camera (Olympus SIS, Japan) and operated at 80 kV.

### Hi-C library preparation

Cells were resuspended in PBS and fixed in 1% paraformaldehyde for 10 minutes with constant mixing. Fixation was quenched by incubating with 125 mM glycine for 10 minutes, followed by centrifugation at 1000×g for 10 minutes at 4 °C and a wash with PBS. The cell pellets were snap-frozen in liquid nitrogen for subsequent processing.

Hi-C libraries were prepared following the Hi-C 2.0 protocol (Belaghzal et al. 2017) with minor modifications. Cells were lysed on ice in a buffer containing 10 mM Tris–HCl (pH 8.0), 10 mM NaCl, 0.5% Igepal CA-630. The pellets were washed twice in NEB 3.1 buffer supplemented with 0.5% SDS. SDS was quenched by incubating with 3% Triton X-100 for 10 minutes. Chromatin was digested with DpnII at 37 °C overnight, and 5′-overhangs were filled using biotinylated dCTP. Ligation of chromatin fragments was performed overnight at 16 °C. Crosslinks were reversed by overnight incubation at 65 °C in the presence of proteinase K. Following DNA extraction, dangling ends were removed with T4 DNA polymerase. The KAPA HyperPlus kit was used for NGS library preparation. Biotin-filled DNA fragments were pulled down using Dynabeads MyOne Streptavidin C1, and the libraries were subsequently amplified.

### Hi-C data processing

Raw Hi-C paired-end NGS data was processed with a modified Juicer (Durand et al. 2016) pipeline (https://github.com/genomech/juicer1.6_compact). It includes several steps, but in the end it returns a merged_nodups.txt file of clean reads, that can be used to make a .cool file using the cooler cload pairs CLI command (Abdennur and Mirny 2020).

### Contacts vs genomic distance

Analysis of change in contact frequency with increasing genomic separation was performed using cooltools (Open2C et al. 2024), particularly the expected_cis() function, which averages values for each individual diagonal on Hi-C matrix at 10kb resolution. Since values on one diagonal are products of contacts over the same distance s, their average can be also seen as interaction frequency for the distance s. Putting together results for a range of distances, a smoothened curve can be plotted, as well as its derivative (using gradient() function from numpy python library). For this analysis, X and Y chromosomes were removed from .cool files, containing Hi-C data, as they are known to have different distribution of contacts. Centromere regions were ignored for the same reason. In order to alleviate the comparison process, final P(s) curves were adjusted to pass through (P(s)=0.1, distance=1*10^5) point by adding [-1- log10(contact frequency over 10^5 bp)] to each individual P(s) value.

For single cell data analysis was consistent with description above, but the demultiplexing stage preceded all the others and was made by scHICExplorer CLI tool (Wolff et al. 2020).

### Compartmentalization. Saddle strength plots

Saddle strength plots were made using cooltools and cool files at 10 kb resolution. After eigendecomposition GC-content was computed for each individual bin, as it helps to properly orient the sign of the first eigenvector (E1). After that, positive values of E1 correspond to the A compartment (active chromatin) and the B compartment (inactive chromatin). After that, regions with similar values of eigenvectors were grouped together, and their interaction frequency with other groups of regions possessing different values of E1 were computed. As a result, a matrix of counts was obtained, values were normalized by dividing each observed value by the corresponding expected value.

### Pileup analysis

Pile-up analysis and insulation score calculation were performed using coolpuppy (10.1093/bioinformatics/btaa073). Submatrices on the intersections of ChIP-seq signals were aggregated, averaged, and normalized (observed over expected). Different sets of parameters were passed into coolpuppy throughout analysis:

Aggregated TAD analysis: (rescale=True, rescale_flank=0.5, rescale_size=99, nproc=16, local=True, min_diag=0). To visualize change between experiments, each individual value was divided by the total matrix. Then one normalized matrix could be subtracted from another.

Aggregated loop analysis: (local=False, min_diag=1, mindist=1000, maxdist=1_000_000, flank = 200_000).

For the pile-up of TADs a consensus estimate of the TAD layout was used (Singh and Berger 2021).

Several pieces of ChIP-seq preprocessed data in a form of bed files from ENCODE were used:

CTCF ChIP-seq for aggregated loop analysis (Mus musculus strain Bruce4 ES-Bruce4): (https://www.encodeproject.org/experiments/ENCSR000CCB/, doi:10.17989/ENCSR000CCB).

### The Rabl structure visualization

Interchromosomal interactions pile-up was performed by a custom script. First, all interchromosomal interaction matrices were extracted from a 100 kb resolution cool file and averaged. The final matrix was visualized as a heatmap. To compare aggregated matrices from different experiments, they were normalized using Knight-Ruiz (KR) matrix balancing algorithm (Knight and Ruiz 2013).

### Bulk RNA-seq

Cells were washed in ice-cold PBS and lysed in TRI Reagent® (Sigma-Aldrich). Total RNA was extracted using the Direct-zol™ RNA Miniprep Plus Kit (Zymo Research), following the manufacturer’s instructions. RNA sequencing was performed by BGI Genomics.

### Bulk RNA-seq data analysis

Raw RNA-seq reads were aligned to the mm10 genome using STAR (2.7.11b) with default parameters, and resulting alignments were quantified using Salmon (1.10.3) to generate a gene expression count matrix. Differential expression analysis was performed using DESeq2 (1.46.0) employing its default settings. Additionally, genes that have less than 5 counts in each experiment were removed from analysis.

## Acknowledgments

Analysis of 3D conformation of the chromatin using Hi-C method was supported by Russian Science Foundation grant #23-74-00055. Cell culture was performed at the Collective Center of ICG SB RAS “Collection of Pluripotent Human and Mammalian Cell Cultures for Biological and Biomedical Research”, project number FWNR-2022-0019 (https://ckp.icgen.ru/cells/). We thank Konstantin Orishchenko group at Novosibirsk State University (supported by the Ministry of Education and Science of the Russian Federation, grant #FSUS-2024-0018) for help with data analysis on computational nodes of Novosibirsk State University. We also acknowledge the Multiple-Access Center for Microscopy of Biological Subjects, ICG SB RAS and Subdiffraction Microscopy and Spectroscopy Core Facility of Moscow State University for granting access to their equipment. The analysis of chromocenter clustering using microscopy methods was carried out with the support of a grant from the state program of the federal territory “Sirius” “Scientific and technological development of the federal territory “Sirius” (Agreement No. 26-03 dated September 27, 2024).

## Author contributions

Conceptualization, A.Y. and N.B.; Investigation, A.Y., E.T., A.S., T.S., I.M., I.P., M.G., E.K., I.K., M.N.; Supervision, V.F. and N.B.; Funding acquisition, V.F. and N.B.; Writing – original draft, A.Y. and N.B.; Writing – review & editing, all authors.

## Data availability

The raw sequencing data have been deposited in the NCBI SRA database with the following accession number PRJNA1300609.

## Supplementary figures

**Figure S1.**
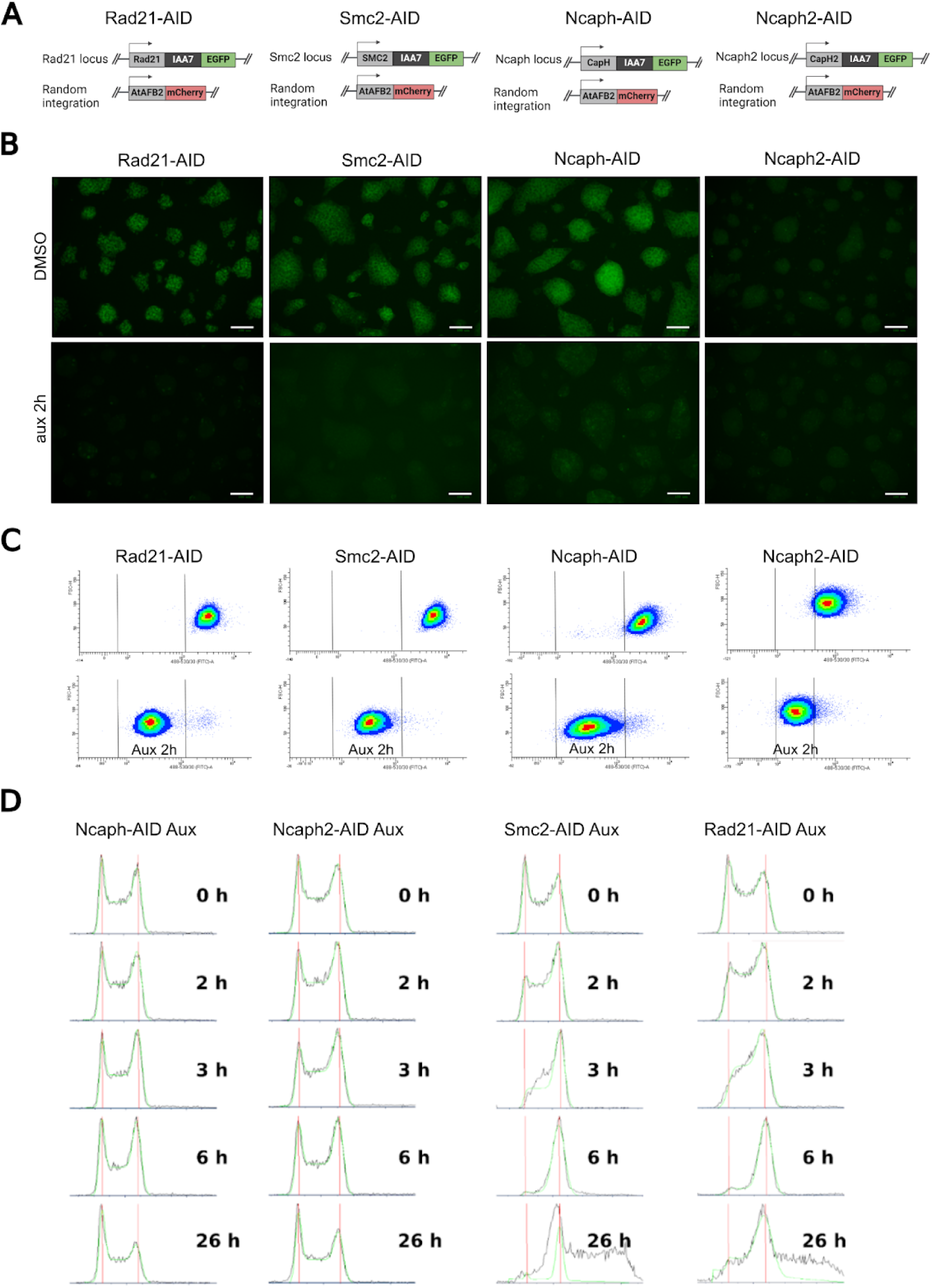
Characteristics of the auxin-inducible degron lines. **A** - A scheme representing the genotype of all degron lines used in this study. **B** - Immunofluorescence microscopy images of the used degron lines exposed to auxin for 2 hours compared to untreated cells. Scale bar 100 μm. **C** - Representative flow plots of the degron lines subjected to 2h treatment with auxin. The vertical lines indicate gates set using GFP-negative cells. **D** - Cell cycle profile of the degron lines following 26h auxin treatment. All samples were prepared in triplicate.

**Figure S2.**
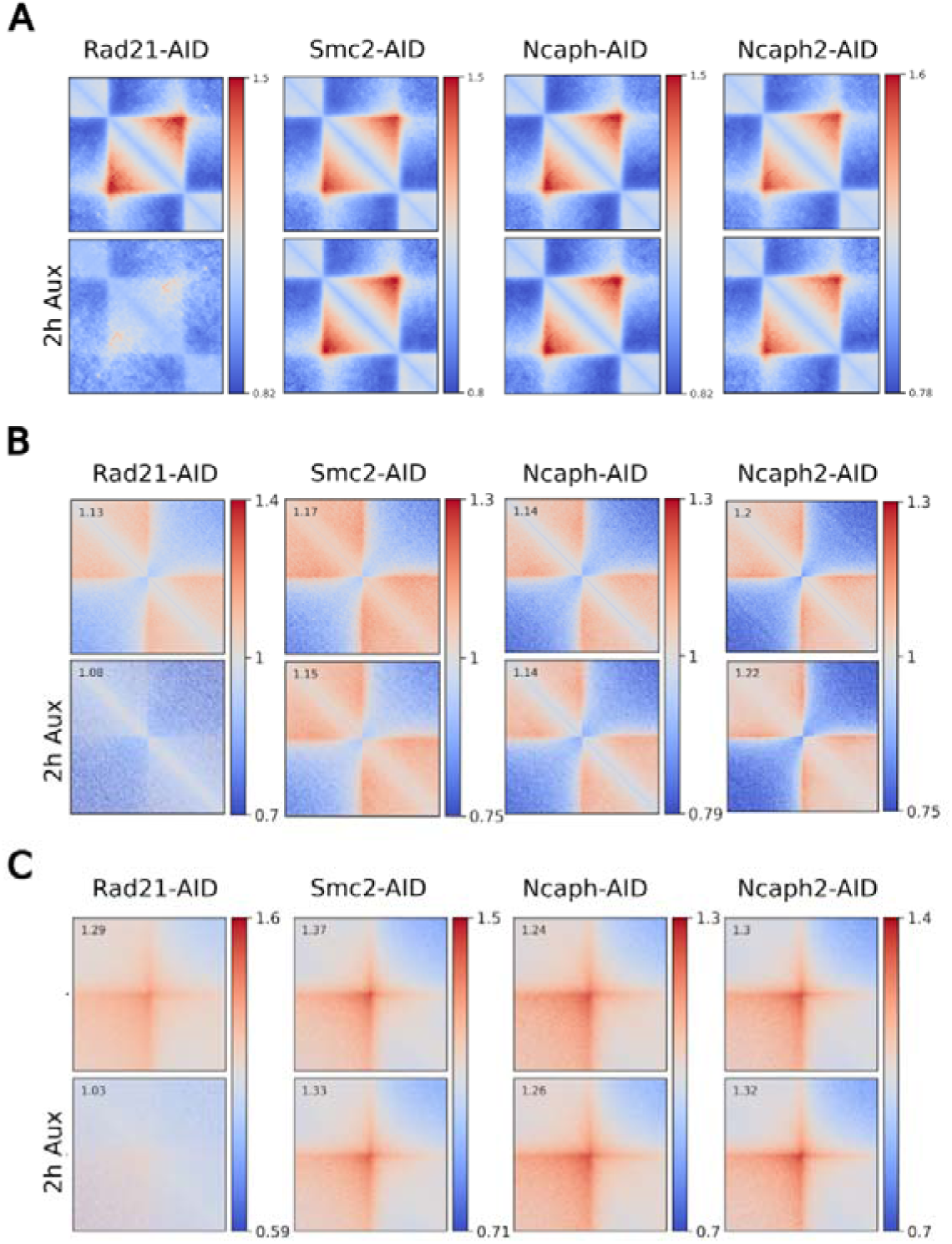
Acute depletion of condensins does not lead to reorganization of interphase chromatin at TAD and loop scale. **A** - Aggregate TAD analysis in cells depleted cohesin (Rad21-AID), condensin I (Ncaph-AID), condensin II (Ncaph2-AID), and both condensins (Smc2-AID) for 2h. **B** - Insulation score of the TAD borders in cells depleted cohesin (Rad21-AID), condensin I (Ncaph-AID), condensin II (Ncaph2-AID), and both condensins (Smc2-AID) for 2h. **C** - Aggregate loop analysis in cells depleted cohesin (Rad21-AID), condensin I (Ncaph-AID), condensin II (Ncaph2-AID), and both condensins (Smc2-AID) for 2 h.

**Figure S3.**
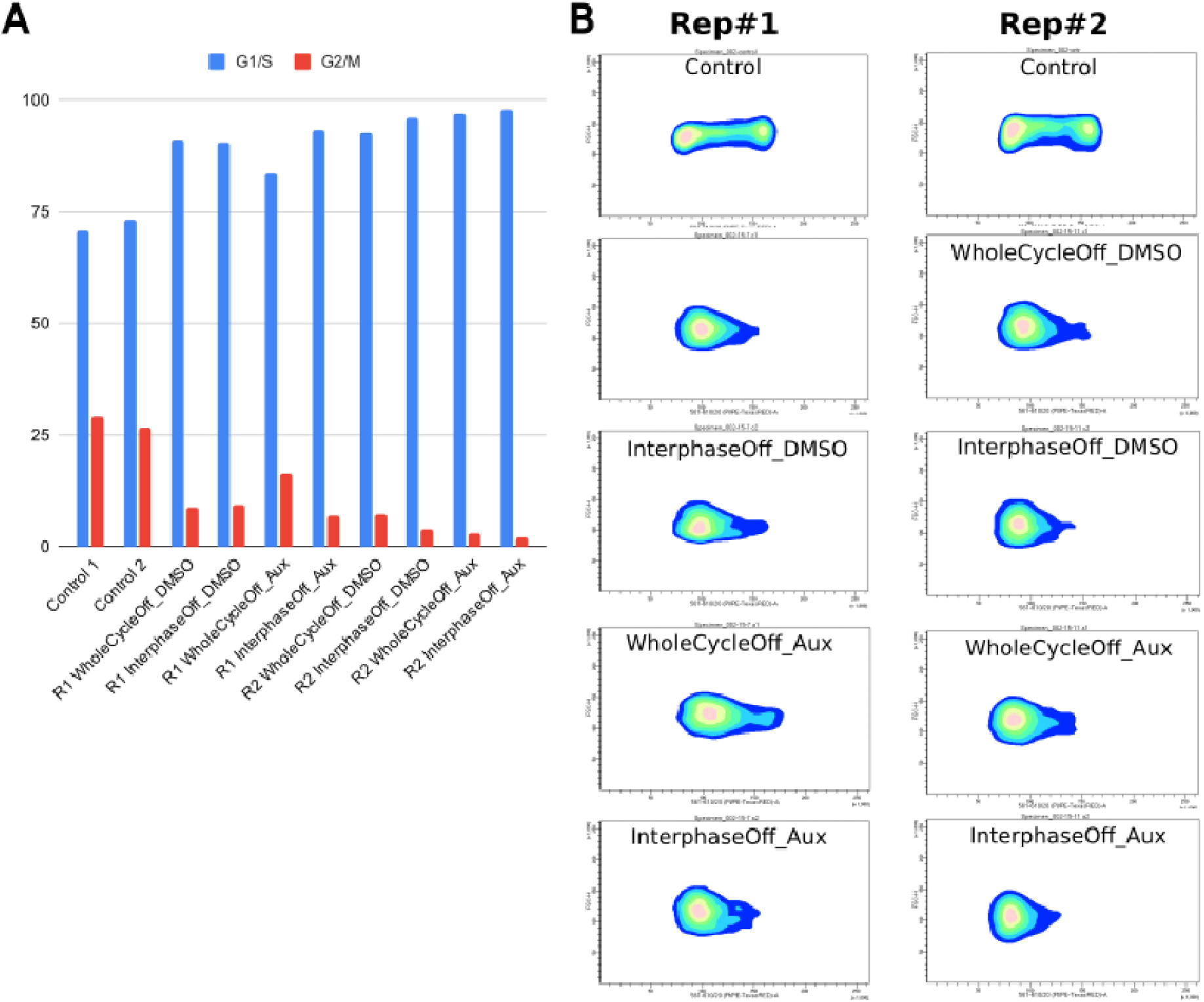
Cell arrest at the G1/S transition with single thymidine blocking. **A** - The percentage of cells at G1/S and G2/M cell cycle stage for replicates correspond to Figure 1E determined by FACS analysis. **B** - Representative FACS images of cells treated with thymidine and auxin correspond to Figure 1E.

**Figure S4.**
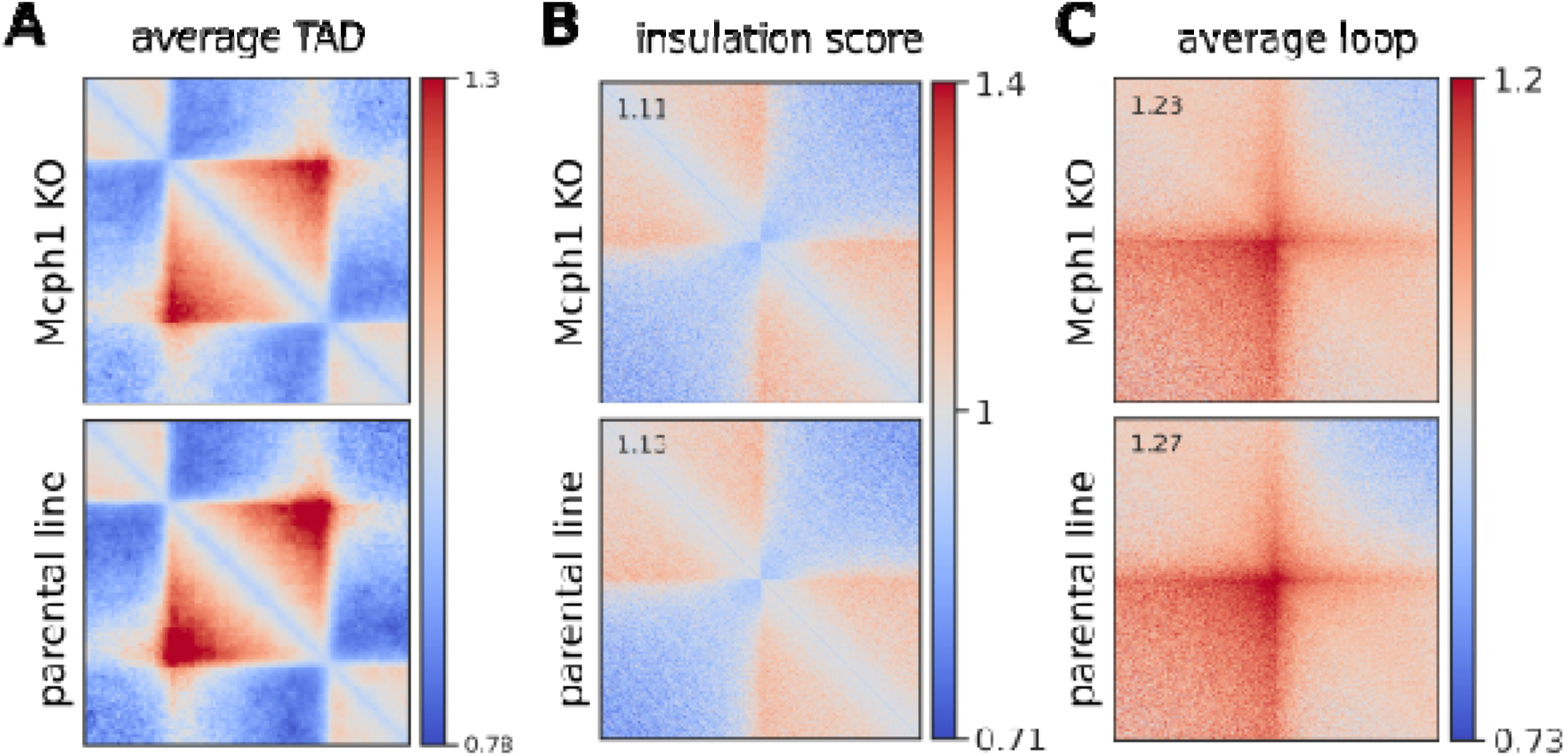
Overloading of condensin II does not affect cohesin-mediated structures such as TADs and loops. (**A-C**) Aggregated TAD (A), insulation score (B) and aggregated loop (C) analysis in Mcph1-KO cells (after downsampling) compared to parental line at 10 kb resolution.

**Figure S5.**
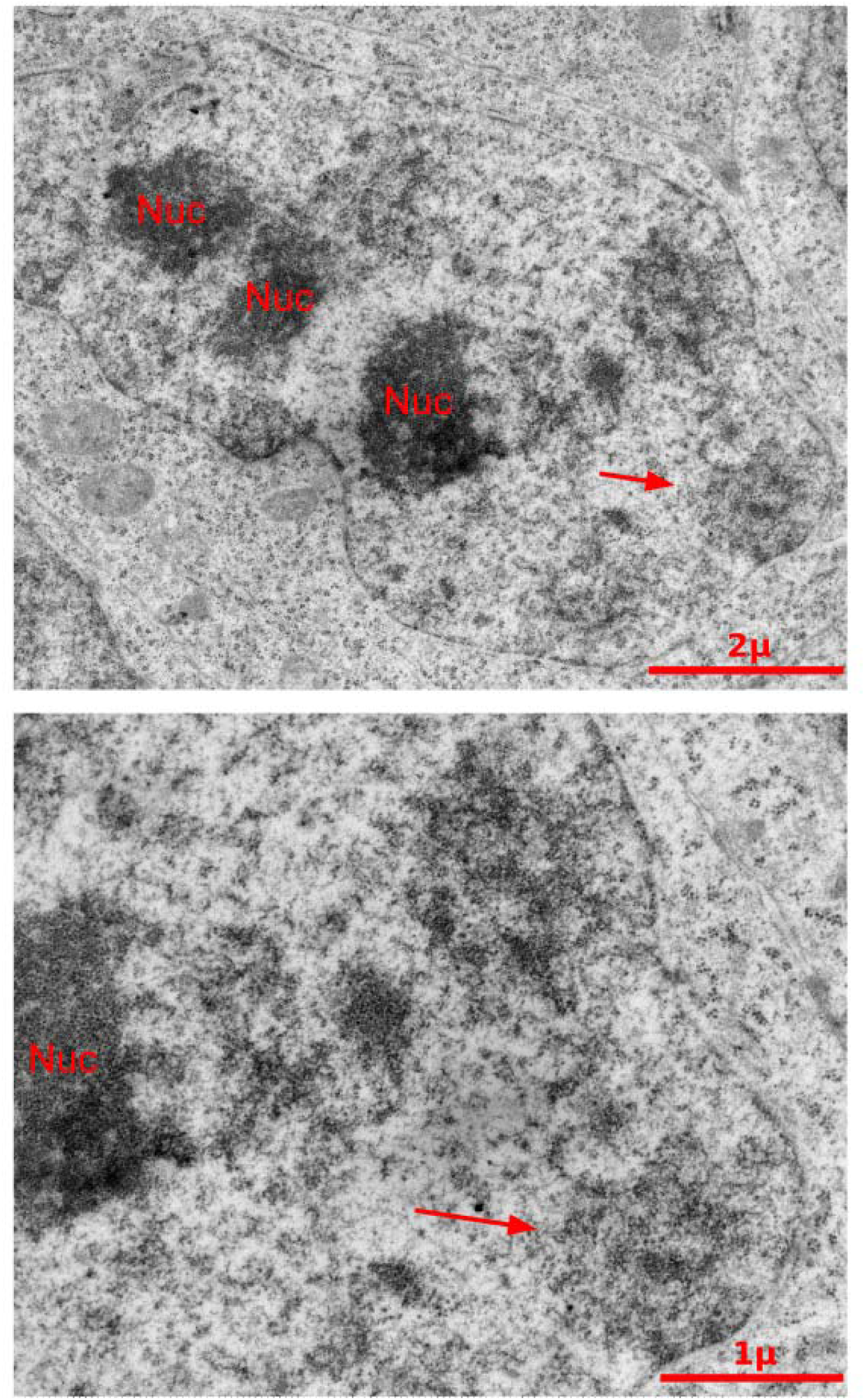
Electron microscopy analysis of prophase-like condensed chromosomes in Mcph1-KO cells. Arrow indicates the heterogeneous areas with a high and low chromatin density.

**Figure S6.**
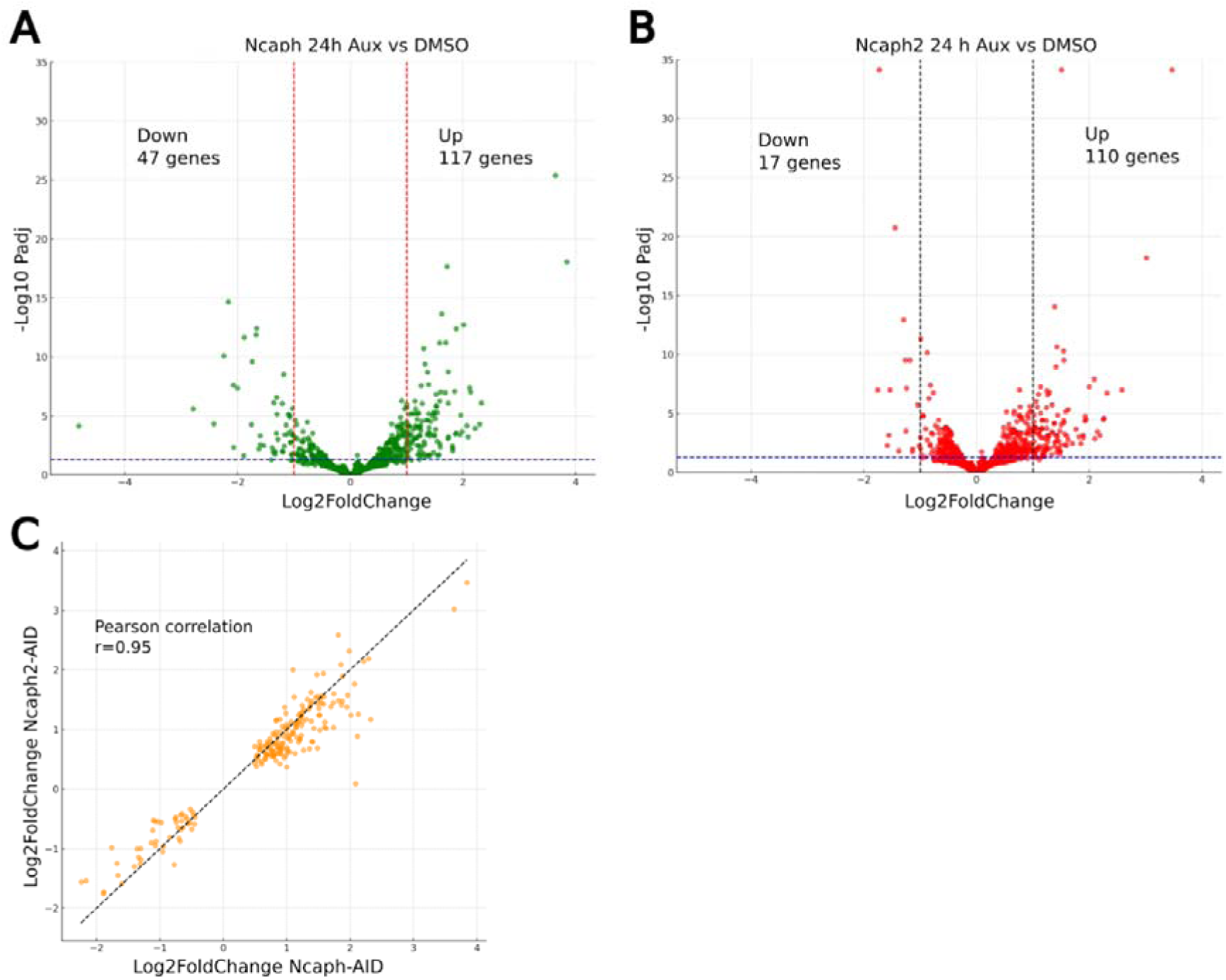
Chromosome configuration changes following chronic condensins depletion do not affect gene expression. (**A, B**) - Volcano plots of the significant DEGs between untreated cells and cells depleted condensin I (A) and condensin II (B) for 24 h. **C** - Pearson correlation between DEG for Ncaph-depleted and Ncaph2-depleted cells.

